# scPanel: A tool for automatic identification of sparse gene panels for generalizable patient classification using scRNA-seq datasets

**DOI:** 10.1101/2024.04.09.588647

**Authors:** Yi Xie, Jianfei Yang, John F Ouyang, Enrico Petretto

## Abstract

Single-cell RNA sequencing (scRNA-seq) technologies can generate transcriptomic profiles at a single-cell resolution in large patient cohorts, facilitating discovery of gene and cellular biomarkers for disease. Yet, when the number of biomarker genes is large the translation to clinical applications is challenging due to prohibitive sequencing costs. Here we introduce scPanel, a computational framework designed to bridge the gap between biomarker discovery and clinical application by identifying a minimal gene panel for patient classification from the cell population(s) most responsive to perturbations (e.g., diseases/drugs). scPanel incorporates a data-driven way to automatically determine the number of selected genes. Patient-level classification is achieved by aggregating the prediction probabilities of cells associated with a patient using the area under the curve score. Application of scPanel on scleroderma and COVID-19 datasets resulted in high patient classification accuracy using a small number (<20) of genes automatically selected from the entire transcriptome. We demonstrate 100% cross-dataset accuracy to predict COVID-19 disease state on an external dataset, illustrating the generalizability of the predicted genes. scPanel outperforms other state-of-the-art gene selection methods for patient classification and can be used to identify small sets of reliable biomarker candidates for clinical translation.

## Background

Single-cell RNA sequencing (scRNA-seq) has emerged as a powerful tool for unraveling cellular heterogeneity within complex biosamples like peripheral blood mononuclear cells (PBMCs)^1^. As scRNA-seq evolves into a mature technology, it is now possible to routinely profile hundreds of thousands of cells from cohorts spanning tens to hundreds of patients^2^. This is further facilitated by the advent of sample multiplexing technologies employing combinatorial barcoding strategies, such as SPLiT-seq^3^. These combinatorial barcoding approaches achieve similar accuracy as commercial droplet-based methods while enabling the profiling of hundreds of samples in a single assay^4^.

These population-level single-cell studies profiling the entire transcriptome presents a unique opportunity for automatic classification of patients’ disease states by learning gene expression patterns specific to the phenotype of interest. However, due to the high sequencing cost of the entire transcriptome, it is challenging to apply single-cell profiling in clinics for each patient sample. Thus, it would be advantageous to identify a small but highly informative gene panel i.e., biomarkers stratifying patient groups, treatment responders vs non-responders^5^ to be used to classify patients. A set of independent patient samples, effectively a test set, is required to validate the predictive power of the classifier and avoid selecting genes specific to certain patients. Moreover, not all cell types have substantial changes in gene expression following perturbation (i.e. diseases, drugs)^6^. Even within the same cell type, the extent of response to perturbations can vary among individual cells. Therefore, it is important to account for the cell differences in contribution to the patient-level classification.

Machine learning approaches have been developed for patient-level classification from scRNA-seq data. CloudPred predicts patient outcomes by modeling the abundance of cell subpopulations using a gaussian mixture model^7^. ProtoCell4P uses prototype-based neural networks to classify patients based on gene expression values^60^. However, both methods use all genes in the patient-level classification, making it prohibitively expensive to deploy in the clinic. Furthermore, both approaches lack gene-level interpretability to explain their prediction. To identify disease biomarkers from scRNA-seq data, several approaches have been developed including ActiveSVM and COMET. ActiveSVM is a machine learning based active forward selection approach where the misclassified cells from the previous iteration are used to select the next most important gene for inclusion into the gene panel^8^. COMET is a statistical method employing a XLLJminimal HyperGeometric test to exhaustively search for gene combinations that can best distinguish a specific cell population^9^. While both approaches identify compact gene panels, they do not provide a way to validate the predictive power of these panels in classifying patient samples. COMET only outputs a list of genes ranked by p-values. ActiveSVM outputs a list of genes ranked by their ability to classify cells but not patients. Furthermore, cells from the same patients are used in both training and testing steps, leading to the potential selection of patient-specific genes during training, which may artificially inflate testing accuracy.

To address these challenges, we developed the scPanel tool, a computational framework to automatically identify a small set of genes for patient-level classification from scRNA-seq data. Specifically, to account for the cell type differences in contribution to patient classification, scPanel first selects cell types responsive to perturbations by quantifying the separation between two patient groups in each cell type. A wrapper-based feature selection method is used to automatically identify a non-redundant minimal gene panel for the selected cell type. The performance of the minimal gene panel is then evaluated by classifying a set of test patient samples, either from the same cohort or a new dataset. Patient-level classification is achieved by aggregating the prediction probabilities of cells associated with a patient using the area under the curve score, thereby accounting for the varying contributions of cells to patient classification. Beyond these challenges, the design of scPanel makes it able to automatically determine the number of selected genes in a data-driven way. It allows easy integration with other methods such as Seurat V3 CCA to remove batch effects from the testing data to enable cross dataset (cross batch) prediction.

To demonstrate the utility of scPanel, we applied it to a diffuse scleroderma patient dataset with a sizable patient cohort^10^ and demonstrated that scPanel can identify a minimal gene set to accurately classify diffuse scleroderma patients. Subsequently, we validated the ability of scPanel-derived biomarkers to generalize to new datasets by applying the tool to two severe COVID-19 PBMC datasets and an infection PBMC dataset with varying COVID-19 severity and influenza patients^11–13^. scPanel-derived biomarkers show consistent superior performance in classifying diffuse scleroderma patients and severe COVID-19 patients when compared with ActiveSVM, COMET and DE genes. The scPanel tool is available at https://test.pypi.org/project/scPanel/ as an open-source python package.

## Results

### Overview of scPanel

We developed a machine learning based tool named scPanel to identify a minimal gene set with predictive power to classify patient-level perturbations (such as disease severity, treatment responses) from scRNA-seq data (Fig. 1, Methods). scPanel comprises three key steps. First, it identifies the most responsive cell type to perturbation. Second, a minimal number of informative genes are then selected from this cell type using a wrapper-based feature selection method^14^. Third, the selected cell types and genes are used to train multiple machine learning models to arrive at a consensus prediction at the patient-level.

**Figure 1.**
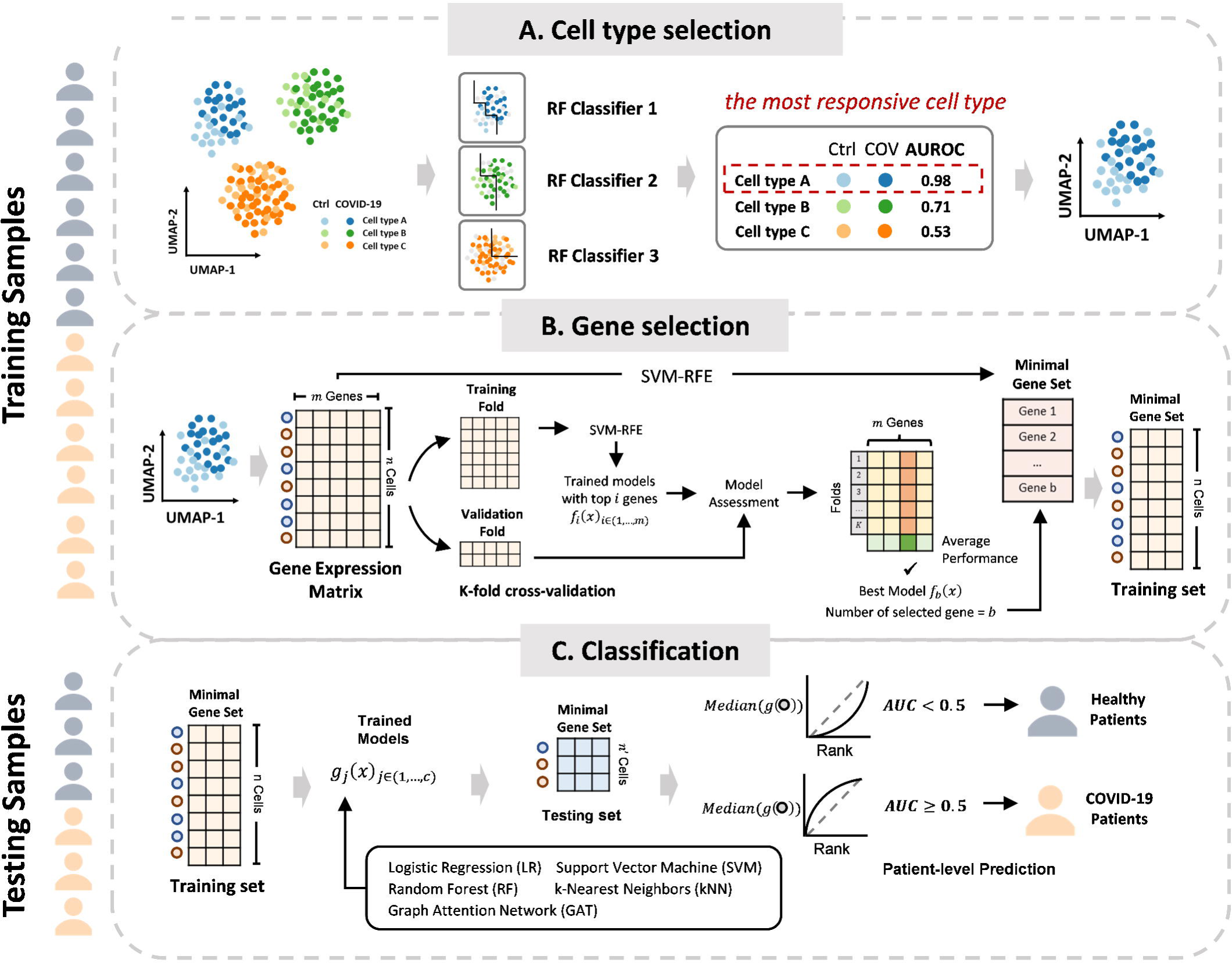
Workflow of the scPanel tool. The scPanel tool comprises three main procedures: **A) Cell Type Selection**: Training samples, annotated with disease status labels and cell types, are divided into internal training and validation datasets. A Random Forest classifier is trained for each cell type and evaluated using the Area Under the Receiver Operating Characteristic (AUROC) score to determine the impact of disease perturbation on each cell type. **B) Gene Selection**: For the selected cell type(s), the minimal gene set is chosen using Support Vector Machine-Recursive Feature Elimination (SVM-RFE) based on k-fold cross-validation to ensure model parsimony. **C) Classification Model Training and Testing**: The training and testing data are refined using the selected cell types and genes. Classification models are then trained, followed by computing the sample-level AUROC score and p-value to predict each patient’s disease status in the testing samples.

The scPanel tool begins with identifying responsive cell types from the entire scRNA-seq dataset. This is because changes of gene expression after perturbation may not occur in all cell types [Perez et al.]. This cell type identification step builds upon the previously established method Augur^15^, which assumes that a cell type that is more responsive to perturbations should be more separable than less responsive counterparts. A random forest (RF) classifier is then trained to predict the perturbation status for each cell type, with the area under the receiver operating characteristic curve (AUROC) score used to quantify prediction performance. A higher AUC indicates a larger separation in a particular cell type, corresponding to greater response to perturbations. However, a significant limitation of Augur is that it assumes that cells from different samples are identical, leading to the training and testing of the cell type-specific RF classifier on cells from the same sample. This assumption can result in the learning of sample-specific patterns during training, thereby inflating testing performance, which is the cell type responsiveness score. scPanel improves upon this by splitting training and testing data by samples, ensuring that the assessment of response to perturbation is generalizable to all patients. This procedure is done repeatedly by sampling an equal number of cells from each patient and giving higher weights to the minority class to reduce biases from dataset compositions.

scPanel constructs minimal gene sets for the selected cell type(s) through SVM-RFE (Support Vector Machine Recursive Feature Elimination), a wrapper-based backward elimination feature selection method^14^. Briefly, all available genes are used initially for classification and the least informative genes are then removed iteratively to search for the best combination of genes. The number of genes we selected is compared against its corresponding classification performance to find the minimal number of genes. This procedure is done in a k-fold cross validation manner to ensure that the number of selected genes is robust across patients. To improve efficiency, gene elimination has been implemented to reduce by percentage (default 3%) at each iteration. Inspired by the findPC tool^16^, the optimal number of genes is automatically determined using the perpendicular line method, which corresponds to the gene count just before a rapid decline in accuracy (See Methods).

To evaluate the effectiveness of the predicted biomarkers, selected cell types and genes are used to train cell-level classifiers with five different machine learning or deep learning algorithms, Logistic Regression (LR), Support Vector Machine (SVM), Random Forest (RF), k-Nearest Neighbors (kNN) and Graph Attention Network (GAT) to account for different complexity within scRNA-seq datasets^17–21^. Cell-level consensus prediction probabilities are summarized by taking the median out of five classifiers. Patient-level classification is determined by aggregating cell prediction probabilities using the area under the curve (AUC) score, with a corresponding p-value to quantify significance (See Methods). In scPanel, all classifiers are weighted by the number of cells in each patient and the number of cells in each class to avoid bias in dataset compositional differences.

### Minimal gene sets identification and patient classification with scPanel

We tested scPanel on four scRNA-seq datasets: a diffuse scleroderma (dSSc) skin dataset (gur2022ssc), two severe COVID-19 PBMC datasets (wilk2020covid, su2020covid) and an infection PBMC datasets (lee2020cov_flu) with mild COVID-19, severe COVID-19 and severe influenza patients (Supplementary Table 1). We tested scPanel in two settings: within-dataset prediction and cross-dataset prediction. In the within-dataset prediction task, gur2022ssc datasets (n=91, 41 control, 50 SSc) were split into a training set (80% of patients) and a testing set (20% of patients) for model training and evaluation respectively. In cross-dataset prediction, we demonstrated the generalizability of scPanel to predict on datasets generated by different labs by testing the performance of a model trained from the wilk2020covid dataset to predict patients in the su2020covid dataset. The specificity of this severe COVID-19 classifier was then determined by making predictions on COVID-19 and influenza patients in the lee2020cov_flu dataset.

Each dataset was first preprocessed to remove low-quality cells and genes. Each cell’s gene expression is normalized by the total UMI counts using standard pipelines (see Methods). Cell types with at least 20 cells detected in at least 3 patients in each class are retained. For each analysis, patients in training and testing sets are not overlapped to ensure that there is no information leakage during cell type selection, gene selection and classification. AUC score is calculated for each patient in the testing set to evaluate the classification performance along with the patient-level accuracy, specificity, precision, sensitivity and F1 score. The parameters used for each model are provided in Supplementary Table 2.

### Minimal gene set for classifying scleroderma patients

To demonstrate scPanel’s scalability to large scRNA-seq datasets with hundreds of patients, we applied the method to a diffuse scleroderma (SSc) skin dataset containing 35,417 cells from 91 donors (41 control, 50 SSc) (Fig. 2A, Supplementary Fig. 1A). Cell type annotations from the original publication, in total 21 cell types, were input to scPanel (Fig. 2B). Different cell types exhibit varying degrees of response to disease perturbation, with *LGR5*-expressing fibroblasts (Fibro_LGR5) showing the strongest response with an AUC score of 0.907 (Fig. 2C). The analysis from scPanel is consistent with the finding in the original publication that the most substantial transcriptional changes in the fibroblast lineage were observed in the Fibro_LGR5 cell type from SSc patients^10^.

**Figure 2.**
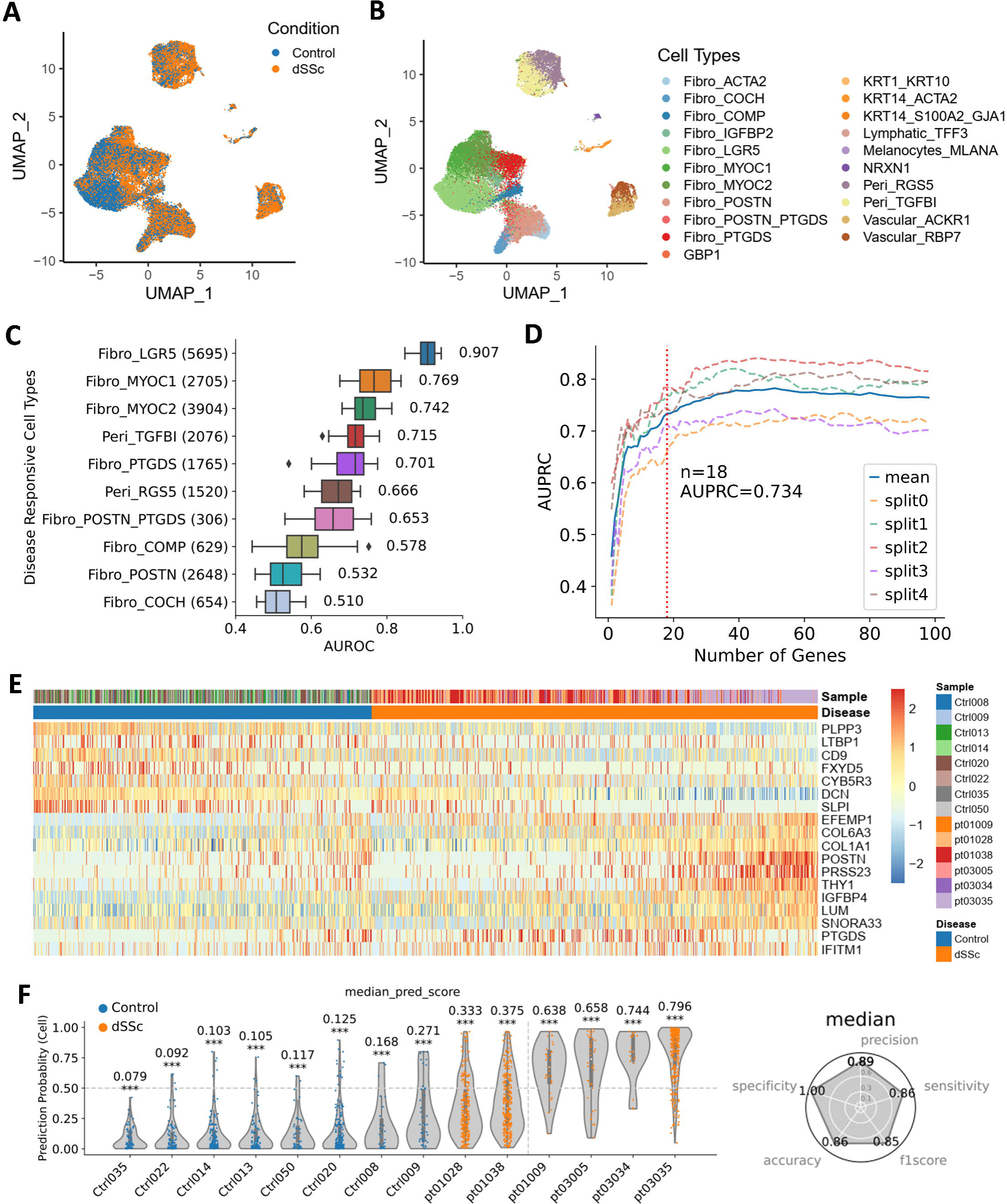
Using scPanel to classify diffuse cutaneous systemic sclerosis (dSSc) patients. A) UMAP of input dataset colored by conditions. B) UMAP of input dataset colored by cell types. C) Cell types ranked by disease responsive score (AUROC). D) Parsimony plot of number of variables (genes) retained and average predictive performance of corresponding model over 5 folds, calculated for Fibro_LGR5 cell type. Dashed line shows the number of genes (n=18) for achieving the parsimonious model. E) Heatmap of gene expression of selected genes in the test set. Values are z-score of normalized gene expression truncated at ±2.5. F) Violin plot of patient-level prediction result. Violins are colored by the label of patients. Y-axis represents cell-level prediction probability summarized by taking the median of 5 classifiers’ prediction probabilities. Value on top of each violin represents sample-level AUC score with p-value (* p<.05; ** p<.01; *** p<.001). The vertical line indicates the patient-level prediction threshold (AUC = 0.5). Samples at the right side of this line are predicted as dSSc patients. Accuracy, precision, sensitivity and recall are computed to evaluate the overall patient classification performance.

We performed gene selection using the Fibro_LGR5 cell type with cells labeled by disease status. 18 genes were identified with 5-fold cross validation, providing enough discriminative power to distinguish dSSc patients (Fig. 2D, 2E). Using these identified genes, Fibro_LGR5-specific SSc classifiers, trained with five machine / deep learning algorithms, achieved high patient-level accuracy (Supplementary Fig. 1D). Median prediction probabilities out of five classifiers are aggregated at patient-level. All patients are accurately classified except for pt01028 and pt01038 with AUC scores of 0.333 and 0.375, respectively. Overall, the patient-level classification accuracy is 0.86 (12 out of 14 patients) in the testing set (Fig. 2F, Supplementary Fig. 1C).

scPanel identified some known and potentially new markers of dSSc within the LGR5-expressing fibroblasts (Fig. 2E, Supplementary Fig. 1B). The *POSTN* gene has been reported to contribute to pathogenesis of scleroderma via *PI3K/Akt* dependent mechanism^22^. The *PLPP3* gene is associated with platelet aggregation in atherosclerosis, a complication common to many SSc patients^23,24^. *PTGDS*, a gene important in prostaglandins (PGs) synthesis, is predictive of poor survival in SSc at protein level^25^. scPanel also identifies multiple genes associated with excessive production and deposition of extracellular matrix (ECM) components (e.g., *COL6A3, COL1A1, LTBP1, EFEMP1, LUM, DCN*) in parallel with the uncontrolled tissue proteases activity (upregulation of the *PRSS23* gene and the downregulation of the *SLPI* gene). Excessive deposition of collagen and ECM proteins as well as the dysregulation of ECM components are known to be the hallmarks of SSc, leading to skin and internal organ fibrosis^26^. The results show that scPanel can automatically identify genes that are involved in SSc pathogenesis and define a minimal gene panel for accurate SSc patients classification.

### Minimal gene set for classifying severe COVID-19 patients

To investigate the wider applicability of scPanel, we applied the method to a severe COVID-19 dataset with 39,196 PBMC cells from 13 patients (6 control, 7 COVID-19)^11^ (Fig. 3A, Supplementary Fig. 2A). Cell type annotations (n = 20 cell types) from the original literature were used as input (Fig. 3B). Most of the cell types have substantial gene expression changes after severe COVID-19 perturbation. CD14+ monocytes showed the strongest immune response with an AUC score 0.999 (Fig. 3C). scPanel showed that six genes from CD14+ monocytes were enough to provide a high accuracy to identify severe COVID-19 patients (Fig. 3D). These six genes (*IFI27, CLU, S100A8, IFITM3, FOS and HLA-DQB1*) showed consistent gene expression patterns between training and testing set (Supplementary Fig. 2B, Fig. 3E). All patients were predicted confidently with AUC scores significantly deviated from 0.5 (random guess) (Supplementary Fig. 2C). This resulted in 100% accuracy (3 out of 3 patients) with AUC scores 0.013 for H1, 0.952 for C3 and 0.977 for C6 (Fig. 3F). All five classifiers consistently showed the same good performance (Supplementary Fig. 2D).

**Figure 3.**
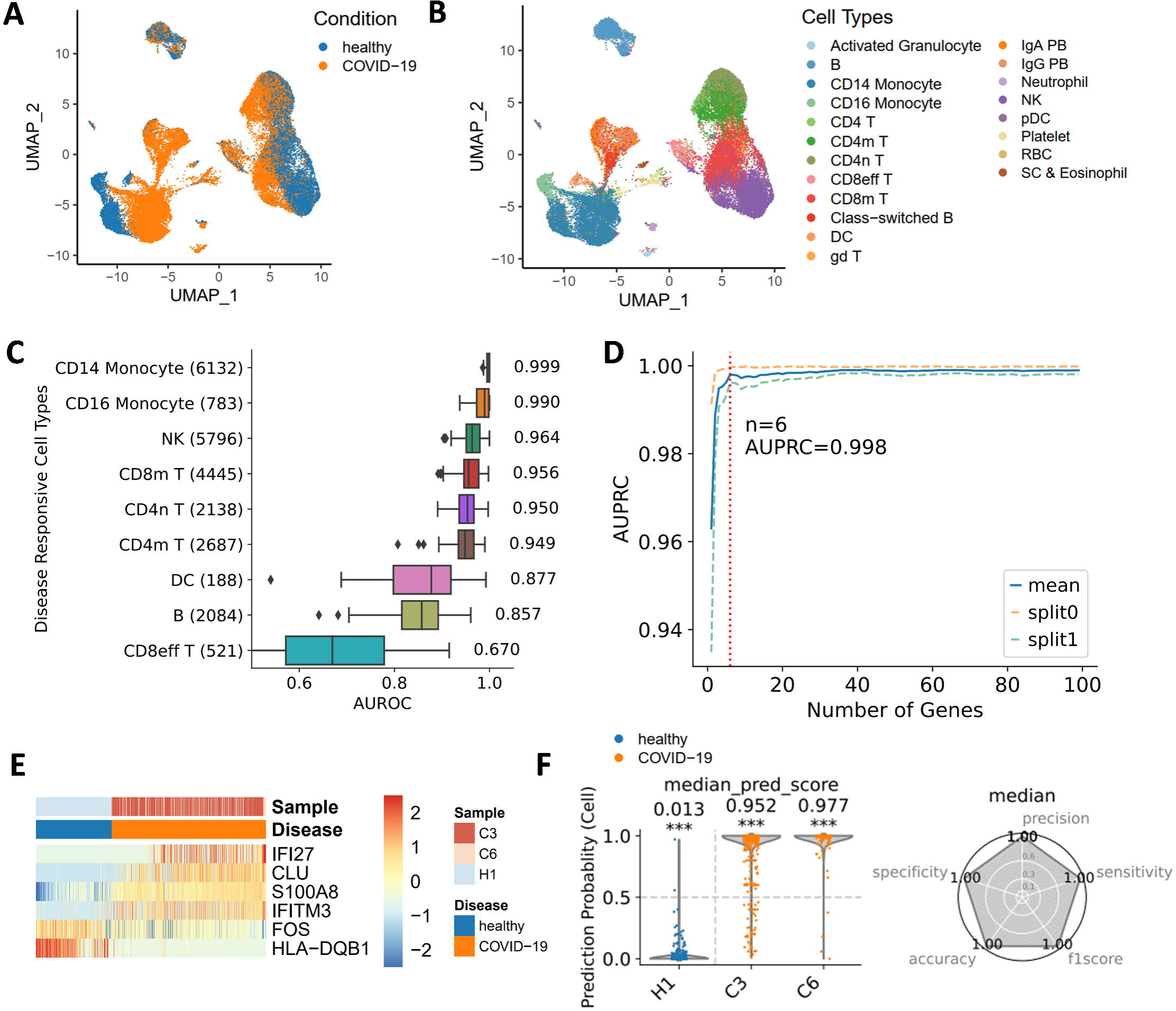

Genes identified by scPanel play important roles in the immune response and inflammatory processes, which are key determinants in the progression of COVID-19. For example, *S100A8* encodes an alarmin protein that mediates host proinflammatory responses during infection^27–29^. It has been shown that *S100A8* is responsible for the induction of an aberrant neutrophil subset in pathogenesis of COVID-19 through stimulating the *TLR4* signal^30^, which are thought to induce cytokine storm or excessive inflammation, as is observed in severe COVID-19 cases^31–33^. *IFITM3* belongs to the interferon-induced transmembrane protein (IFITM) family that is known to play a significant role in antiviral response by restricting virus entry into cells^34–41^. It has been recently reported that SARS-CoV-2 virus hijacks IFITM proteins for efficient infection, which explains the rapid spread of this virus^42^.

### Generalizability of prediction of severe COVID-19 patients

To demonstrate the generalizability of scPanel across different datasets, we applied the severe COVID-19 classifiers to the independent su2020covid dataset (Supplementary Fig. 3A, 3B). Specifically, CD14+ monocytes were computationally isolated from the su2020covid dataset in concordance with the cell type used in the training data. We observed pronounced batch effects between the training set (wilk2020covid) and the testing set (su2020covid) with a clear separation of cells by their dataset-of-origin (Fig. 4A). Consequently, the classifier exhibited a suboptimal performance with an accuracy of 0.67, misclassifying all healthy patients (Fig. 4B).

**Figure 4.**
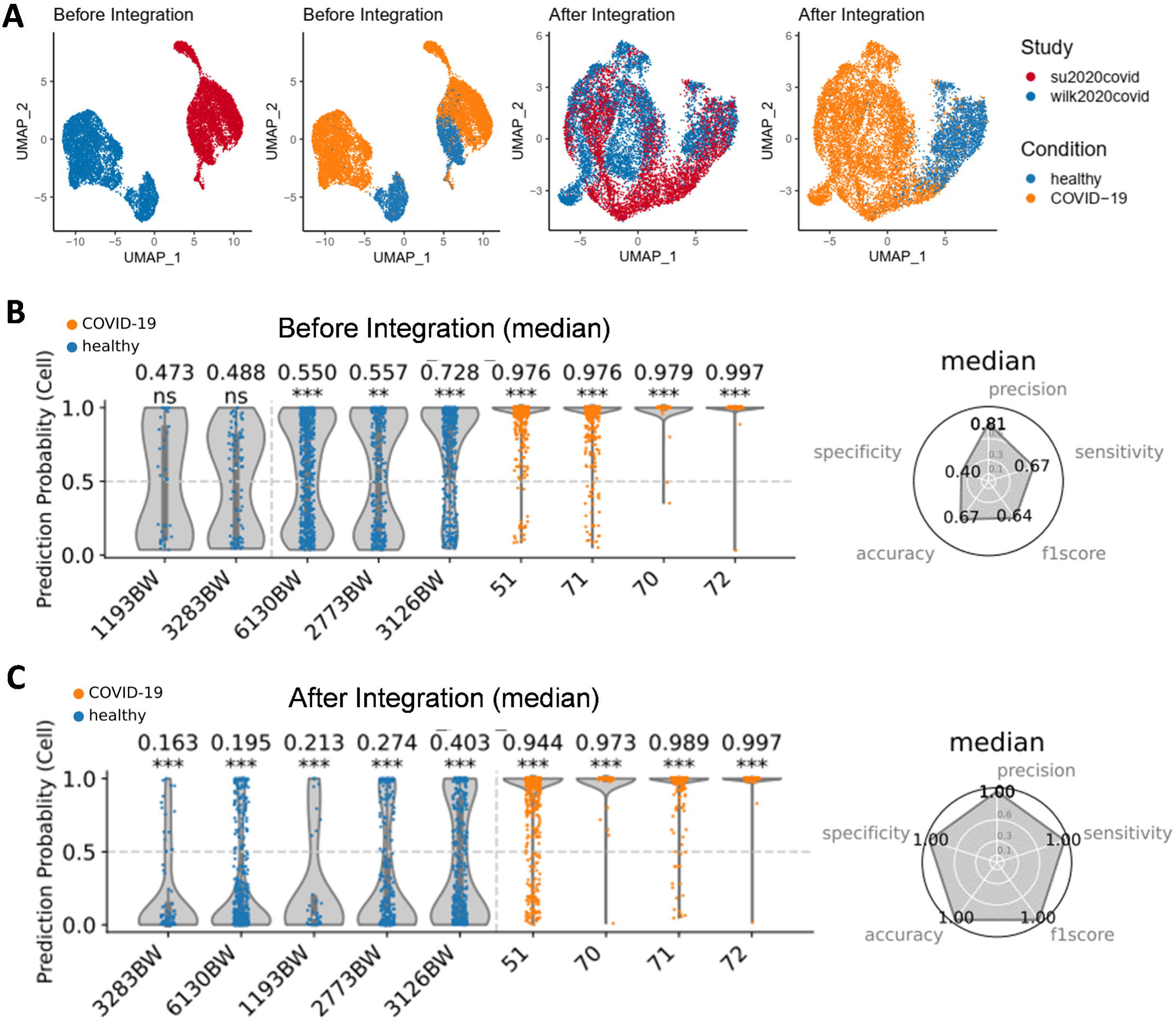

We hypothesized that these batch effects need to be removed for accurate prediction. To address this, we applied the Seurat V3 canonical correlation analysis (CCA) method to remove batch effects for better cross-dataset prediction performance^43^. After removing batch effects from the testing set, we observed a more integrated representation of CD14+ monocytes from both datasets, with a predominant clustering of cells by disease status rather than dataset-of-origin (Fig. 4A). This procedure substantially improved the cross-dataset prediction accuracy, elevating it from 0.67 to 1.00 (Fig. 4C). Interestingly, the GAT classifier showed an inherent robustness to batch effects, consistently maintaining 100% accuracy, irrespective of integration (Supplementary Fig. 3C, 3D). Our findings show that scPanel, providing appropriate batch effect removal, can serve as a tool for patient-level classification in cross-dataset prediction settings.

### Specificity of prediction of severe COVID-19 patients

To assess the specificity of our severe COVID-19 classifiers (LR, RF, SVM, KNN, and GAT), we incorporated a PBMC dataset, referred to as lee2020cov_flu, from patients with various COVID-19 severities and influenza cases (Supplementary Fig. 4A, 4B and 4C). If the severe COVID-19 gene set, along with the trained classifiers, specifically captures the unique expression pattern distinct from that of mild COVID-19 and influenza cases, the classifiers trained on healthy and severe COVID-19 data should yield prediction probabilities randomly distributed between 0 and 1 when predicting moderate COVID-19 and influenza cases. To obtain the prediction probabilities on lee2020cov_flu data, we first integrated it with our training set (wilk2020covid) to mitigate the batch effects (Fig. 5A). After integration, all five classifiers demonstrated a 100% accuracy in predicting healthy individuals and severe COVID-19 patients within the lee2020cov_flu dataset (Supplementary Fig. 4D and 4E). However, LR, RF, SVM and KNN classifiers demonstrated a lack of specificity for severe COVID-19, incorrectly classifying all severe influenza cases as severe COVID-19 with 100% prediction probabilities (Supplementary Fig. 4E). In contrast, the GAT classifier can distinguish the minor difference in gene expression changes between severe COVID-19 and other conditions, assigning a median sample prediction probability of 0.65 to severe influenza cases and 0.45 to moderate COVID-19 cases (Fig. 5B and 5C).

**Figure 5.**
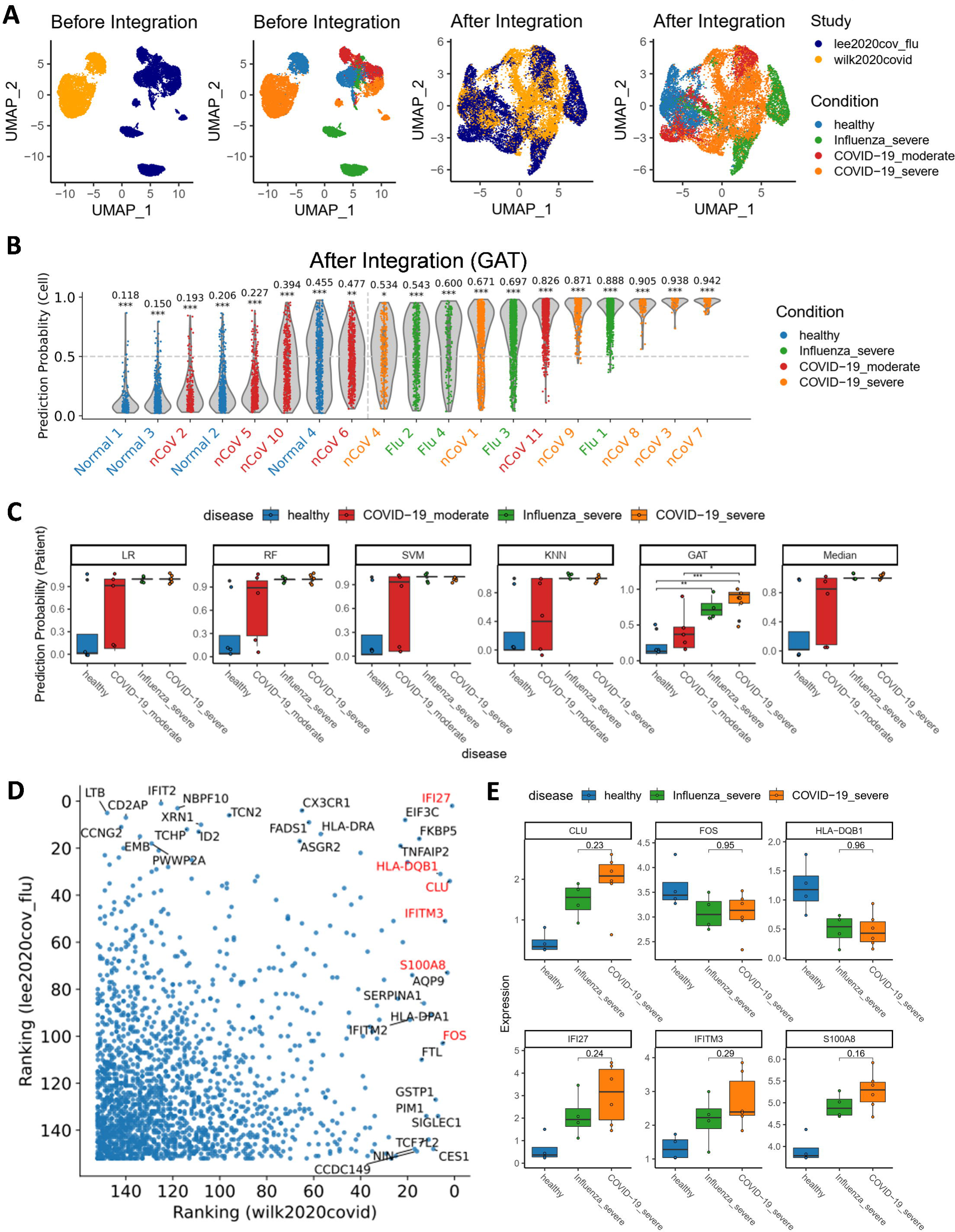

To investigate the potential non-specificity of the gene set used in the severe COVID-19 classifier, we derived a severe influenza gene set from lee2020cov_flu by applying scPanel to healthy and severe influenza groups. We observed that certain genes in the severe COVID-19 gene set, such as *IFI27, HLA-DQB1, and CLU*, also hold significance in the context of influenza patient classification (Fig. 5D). Notably, *IFI27* has been previously described as a biomarker for various infectious diseases including COVID-19 and influenza^44,45^. Furthermore, the expression levels of these severe COVID-19 genes exhibited no significant differences between severe influenza and COVID-19 groups (Fig. 5E).

### Stability analysis

In the default scPanel gene selection procedure, selecting genes from a single training set might lead to overfitting, where the gene expressions describe patterns specific to the training set but do not generalize to other patients. To improve the robustness of our gene selection methodology, we have implemented a stable gene selection mode in scPanel. This enhanced approach involves iterative down-sampling of the training data into multiple subsets, followed by the application of the scPanel gene selection algorithm to each subset. This down-sampling simulates the presence of multiple random cohorts, allowing for the identification of “stable genes” that are consistently being selected across cohorts and thereby reducing the model variance. Genes that are consistently selected in over a fraction (e.g., 50%) of these subsets are retained, culminating in a final, more reliable gene panel (Fig. 6A).

**Figure 6.**
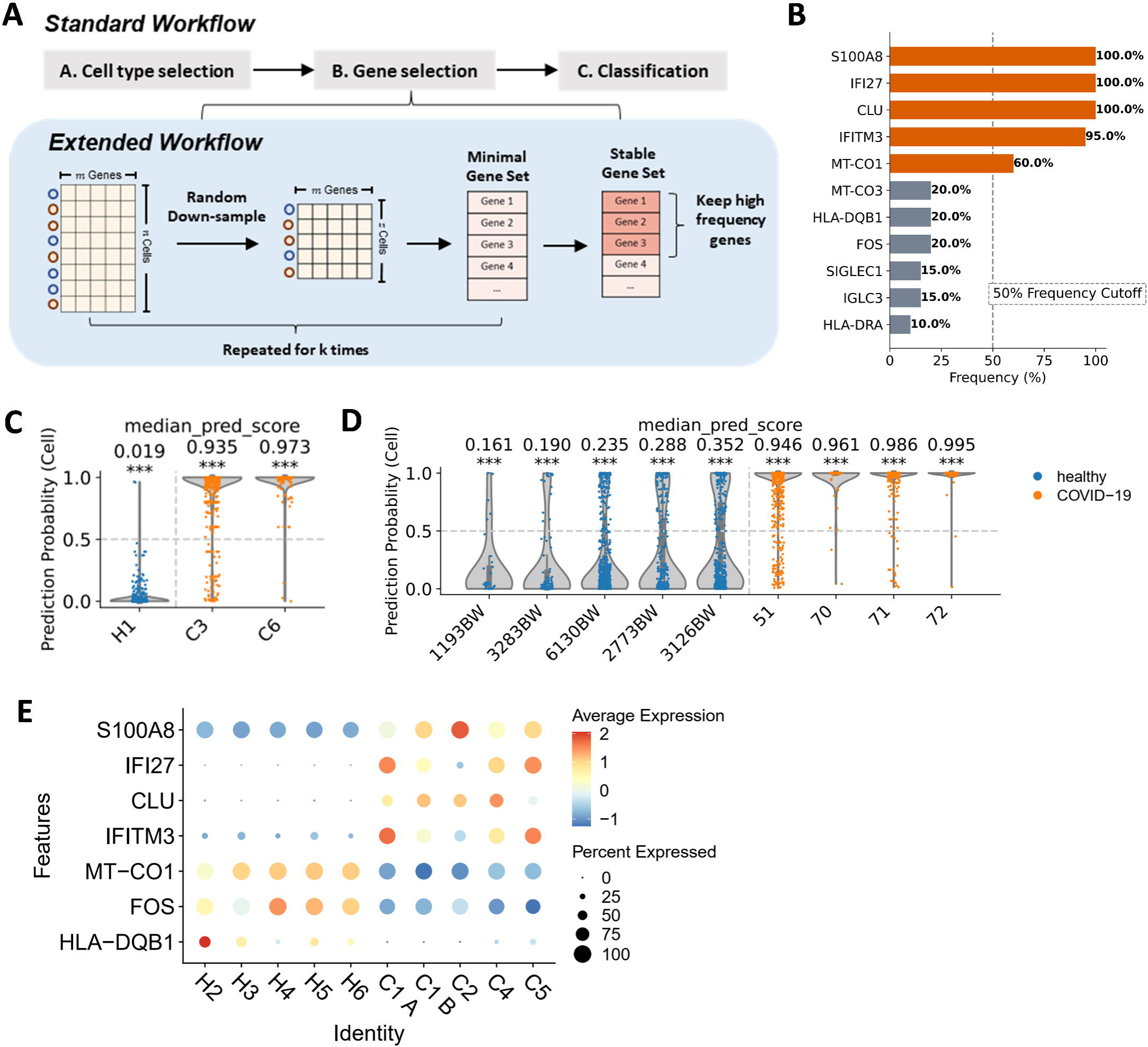

Using this approach, genes *S100A8, IFI27, CLU, IFITM3,* and *MT-CO1* have been identified as the stable gene panel for the classification of severe COVID-19 patients, utilizing the same training dataset as previously employed (Fig. 6B). Classifiers developed using this stable gene panel (comprising five genes) demonstrated comparable performance to that trained with the standard procedure (comprising seven genes), achieving 100% accuracy in both within-dataset and cross-dataset predictions (Fig. 6C and 6D).

Notably, genes like *FOS* and *HLA-DQB1*, which were selected in the standard procedure, were not chosen in the stable mode (Fig. 3E and 6B). Instead, *MT-CO1* was selected more frequently, forming the stable gene set alongside other genes. This preference can be attributed to the fact that *HLA-DQB1* is expressed in a small proportion of cells in the training data (Fig. 6E). Thus, its predictive power gets reduced after data down-sampling. In contrast, MT-CO1, with its stable expression across samples, emerges as a more suitable candidate in the stable gene selection mode (Fig. 6E). A stable gene selection mode has been included in our scPanel python package.

### Benchmarking, Power Analysis and Computational Requirements

To evaluate scPanel’s performance in identifying minimal gene panels, we compared it to several marker identification methods including the traditional differential expressed (DE) gene method, or ActiveSVM and COMET. In dSSc patients classification, scPanel selected 18 genes as the minimal gene panel. Correspondingly, we selected the top 18 genes from the DE gene method and ActiveSVM for comparison. Given COMET’s inherent limitation of only exhaustively searching a maximum of four genes, we additionally conducted a comparison using the top four genes from scPanel. We observed that scPanel consistently outperformed the other methods in classifying ssc patients, both with gene panels comprising either the top four or 18 genes (Table 1). Furthermore, the same benchmarking experiments were conducted in severe COVID-19 patient classification. scPanel maintained its superior performance over the other methods (Table 2), further validating its effectiveness in identifying minimal yet potent gene panels for patient-level classifications.

To assess scPanel’s performance against variations in cell number and sample size, we conducted an empirical power analysis by downsampling the training data to different sizes. In the severe COVID-19 dataset (wilk2020covid), cell-level analysis consistently identified CD14+ monocytes as the most responsive cell type across all subsets (Supplementary Figure. 5A), with some variation in the identified genes (Supplementary Figure. 5B) but a smaller set of stably selected genes (Supplementary Figure. 5C). Despite this, predictions for all patients in the test set remained accurate across all subsets, albeit with a marginal reduction in AUC scores for subsets with size smaller than 70% (Supplementary Figure. 5D). Further cell-level power analysis within the dSSc dataset (gur2022ssc) identified LGR5+ fibroblasts as the most responsive cell type when more than 13,141 cells (60% of the total) were used (Supplementary Figure 6A). Above this number of cells, there are no significant differences in the number of selected genes with similar sets of genes being identified consistently (Kruskal-Wallis test, p=0.18) and patient-level prediction remains the same for LGR5+ fibroblasts (Supplementary Figure. 6B - 6D).

For sample-level power analysis, the dSSc dataset (gur2022ssc) was selected due to its large patient cohort. It was observed that the cell type responsiveness score (AUC) for LGR5+ fibroblasts began to plateau at 40% (33 samples) of the full sample size (Supplementary Figure. 7A). Correspondingly, the gene selection started to exhibit large variations when the sample sizes were smaller than 40% (Supplementary Figure 7B, 7C). Despite the variability in gene selection, the top 5 most important genes (DCN, PLPP3, EFEMP1, IGFBP4 and POSTN) are selected with frequency at least 65% and patient-level prediction remains stable across all subsets with size ≥ 40% (Supplementary Figure 7D). Computationally, both peak memory usage and processing time displayed an almost linear increase relative to different sizes of the input cells and samples (Supplementary Figure. 5E, F, 6E, F, and 7E, F).

## Discussion

In this study, we present scPanel, a computational framework designed with three core functions: 1) identifying cell types responsive to perturbations; 2) uncovering a minimal set of genes predictive of these perturbations; and 3) enabling patient-level perturbation prediction in large-scale single-cell RNA sequencing (scRNA-seq) datasets.

Recently, other methods, namely COMET and ActiveSVM, have been proposed to identify disease specific gene sets and predict patient disease states. However, these methods use all cells in disease states as input, hence making an assumption that all diseased cells are homogeneous as well as the response to a similar extent under perturbation. This overlooks the fact that in some biological settings, only certain cell types may respond to perturbations. For example, in anti-PD1 treatment, only specific macrophage phenotypes (CCR2+ or MMP9+) are correlated with T cell expansion, an indicator of anti-PD1 treatment response, in pre-treatment tissues^46^. Thus, applying a uniform label to all cells, whether they are responding or non-responding counterparts, would dilute the biological signal originating from the responding macrophage subtypes. This makes it difficult for the model to converge and learn gene expression patterns specific to responding cell types. Recognizing the importance of accurately labelling responsive cells in diseased samples in the supervised learning process, scPanel emphasizes prioritizing the most responsive cell type for model training. Uniquely, scPanel addresses patient-level heterogeneity by training on one patient cohort and validating cell type responsiveness and gene importance scores in an unseen patient cohort. This approach ensured that the patient classification model by scPanel, trained with selected cell types and genes, is generalizable across different patients. To accommodate varying data complexities, five distinct algorithms were employed to develop the patient classification model, enhancing its robustness and applicability.

Transcriptomics data frequently exhibit collinearity, enabling the construction of gene modules comprising genes with similar expression patterns^48,61^. Consequently, a small number of informative genes can be extracted to efficiently represent these modules and hence reveal a cell’s state.–scPanel utilized SVM-RFE, a feature selection method widely applied in different omics data ^50–52^, to extract a minimal set of highly informative genes to distinguish perturbations. The selection process is coupled with a cell-state classification task to make sure selected genes have enough predictive power. Unlike other representation reduction methods such as PCA or random composite measurements, scPanel returns interpretable genes instead of complex features such as linear combination of genes^49^. If applied as a clinical tool, the predicted genes can be measured directly without developing complicated scoring metrics.

A well-documented challenge with scRNA-seq data is the prevalence of batch effects^53,54^, where classifiers trained on one batch significantly underperform on a separate, independently collected cohort due to technical disparities in sample collection, reagents, or sequencing platforms. This issue can be addressed by integrating the testing data with the training data. To avoid retraining the model with each new batch of testing data, it is important that the training data remains unchanged during data integration. To solve this, scPanel adopts the Seurat3 CCA method, an approach that integrates data without altering the values in the training data, and one that has demonstrated superior integration performance in recent benchmark studies^55^. Notably, our observations reveal that the GAT classifier, a graph based deep learning model, can predict well on different batches of data, despite being trained only on one batch. GAT, by design, leverages attention mechanisms to model relationships and dependencies between cells to output a more meaningful representation that can be generalized to different batches of data. The other four classification algorithms treat cells independently and find the optimal hyperplane that separates the different classes based on the original data, lacking the capacity to learn a more general representation as GAT.

### Limitations

While scPanel provides a valuable framework for identifying minimal gene panels and classifying patients, it has several limitations. A clinically applicable gene panel ideally exhibits significant expression differences between groups, ensuring detectable signals by technologies like qPCR. Thus, incorporating the magnitude of gene expression differences into the gene selection process could enhance the panel’s discriminating power. Additionally, the high costs of scRNA-seq experiments often limit sample sizes in the range of tens to hundreds of samples, which might not comprehensively represent the entire population. Thus, the generalizability of gene panels derived from scPanel is largely limited by the input training data, which will eventually improve with the emergence of population-level single-cell studies encompassing more patients. Also, the selected gene panel may be predictive of diseases with similar mechanisms, for example as shown in the results where the COVID-19 gene panel also predicts influenza cases. Thus, comparative analyses using similar disease class single-cell data are necessary to further improve the specificity of the identified genes.

Along with its applicability to scRNA-seq data, scPanel framework can be expanded to other data types by merely changing the input. It has potential applications in identifying biomarkers from CyTOF (Cytometry by time of flight)^56^ and other multi-omics single-cell data. Furthermore, when applied to a disease single-cell dataset e.g. COVID-19, scPanel can prioritize candidate driver genes to be perturbed in downstream Perturb-seq experiments^57^. Beyond clinical application, scPanel can prioritize research interests, such as focusing on specific cell types or evaluating the importance of user-defined gene sets.

Although scPanel currently focuses on binary classification, its future expansion to multi-class problems is feasible, given the availability of multi-class SVM-RFE for gene selection^58^. Integrating additional information like cell type abundance and patient-level electronic health records (EHR) could further refine prediction accuracy. We anticipate that scPanel will significantly contribute to enabling the extraction of biological insights in the form of gene panels from single-cell level data for clinical applications.

### Key Points

- We proposed a machine learning based framework scPanel for patient-level classification from scRNA-seq data
- scPanel can identify responsive cell types and a minimal gene set for classification
- scPanel achieved high accuracy with small number of highly informative genes in SSc and COVID-19 classification
- Compared with existing approaches e.g. COMET, ActiveSVM and differential expressed genes, genes selected with scPanel have more predictive power
- scPanel is available as an open-source python package and implemented with parallel computing

## Methods

### Design and implementation of scPanel

scPanel expects an input of normalized single cell gene expression matrix *D*={*x_h,i_*}*_nxK_* where *x_h,i_* is the expression of gene *h* in cell *i*. *n* denotes the number of measured genes and *K* denotes the number of cells. Each cell *i* must have a sample label *Z_i_* indicating which sample this cell belongs to. Cell state labels *Y*={*y*_1_, *y*_2_,. . .*y_i_*}, *y_i_* ∈ {0,1} are required and can be derived from the state of the sample the cells belong to (e.g. cells from COVID-19 patients are labeled as 1 and from healthy controls are labeled as 0). Importantly, it also expects that an initial cell type annotation of cells (denoted as *c*) is available. We denote the cell type labels as *c_i_* = *t,* where *t* ∈ {1, . . . *T*} and *T* is that total number of cell types in *D*.

Data is splitted into training set *D_train_*, *Y_train_* and testing set *D_test_, Y_test_*, with the division based on sample labels and stratification according to cell state labels. Z-score transformation is performed to standardize gene expression. To reduce the impact of technical outliers, the standardized values whose magnitude exceeds 10 are clipped. Gene expression mean and variance of the training data are used to standardize the testing data.

#### Cell type selection

For each cell type, samples with less than 20 cells (default) are removed. Cell types with less than 3 patients in at least one condition are removed. Responsive cells should become more separable from cells in control groups. scPanel qualifies responsiveness of cell types to perturbation by asking how readily the sample labels associated with each cell (for example, treatment versus control) can be predicted from the gene expression. In practice, we treat it as a classification problem. If cells in the treatment group are not responsive while being labeled as positive, the model will not converge during training. Thus, it will have low prediction performance during testing.

For each cell type at each iteration, we sampled (without replacement) 20 cells from each sample. 60% of samples are randomly splitted into the training set. Sample labels (for example, treatment versus control) are used as class labels to train a random forest classifier using scikit-learn with class_weight = ‘balance’ to avoid bias from different numbers of cells from different classes^59^. Classification performance of the model is evaluated on the remaining 40% samples and AUROC is calculated as the metrics to determine the responsiveness of cell types. This procedure is repeated 100 times by default for robust assessment. A Shapiro–Wilk test is carried out to test the normality of the distribution of AUROC values. If the p-value is larger than 0.05, the mean value is used to represent final AUROC for the cell type. Otherwise, the median value is used.

#### Minimal gene panel selection

SVM-RFE was introduced by Guyon et. al., for selecting important genes from gene expression data for cancer classification^14^. SVM-RFE, starting with all the genes, removes the gene that is least significant for classification recursively in a backward elimination manner. In each iteration, the ranking score of genes *s* is computed from the coefficients of the weight vector *w* of a linear SVM as follows:

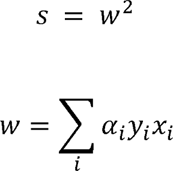

Where ∈ is the class label of the cell *x_i_* and the summation is taken over all the training samples. *α_i_* are the Lagrange multipliers involved in maximizing the margin of separation of the classes. For computational efficiency, 3% of genes are removed at each iteration by default in scPanel. We allow users to adjust this value or remove one gene at a time by inputting different parameters.

The procedures of SVM-RFE are implemented as follows:

1. Start: ranked gene set *R*=[]; selected gene subset *S* = [1,…n];
2. Repeat until all genes are ranked:

a. Given a training set {*x*_1_, *x*_2_,…,*x_i_*, …,*x_l_*} and class labels {*y*_1_, *y*_2_,…,*y_i_*, …,*y_l_*}. Train a linear SVM with genes in set *S* as input variables by minimizing:

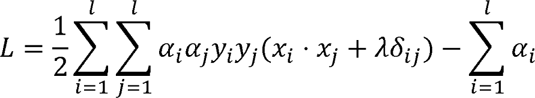 *δ_ij_* is the Kronecker symbol (if *i*=*j* and 0 otherwise), and *α*={*α*_1_, *α*_2_,…*α_l_*} are the parameters to be determined. *Y* and *c* are positive constants for regularization (soft margin parameters).
b. Compute the weight vector;
c. Compute the ranking scores for genes in set S: *s_i_*=(*w_i_*)^2^;
d. Find 0.03 × *n*. bottom-ranked genes *e*=[1,…,0.03 × *n*];
e. Update: *R*=[*e*,*R*], *S*=*S* – *e*;
3. Output: Ranked feature list *R*.

To determine how many top-ranked genes in *R* should be selected from the training data, we adopted SVM-RFE with a N-fold cross validation strategy (SVM-RFECV), where the training data are further partitioned into N equal-sized subsets (folds), N-1 folds are taken as the internal training set and the remaining one fold is taken as the internal validation set. At each fold, whenever the bottom-ranked gene was removed by SVM-RFE, the resulting gene subset was evaluated by summarizing the classification performance of the same SVM model on the internal validation set using AUPRC scores. The minimal number of genes is determined on which the SVM can get optimal AUPRC scores across N folds. The optimal AUPRC scores are determined in a data-driven way by applying perpendicular line method^16^ on the parsimony plot where x-axis is the number of top-ranked genes retained in SVM-RFE in the internal training sets and y-axis represents the average classification performance of the same SVM model in the internal testing sets. scPanel automatically chooses the number of top-ranked genes *M* with the longest perpendicular line to the line passing the two points of the highest and lowest ranks. Top *M* genes from ranked feature list *R* are taken as the minimal gene panel.

To avoid bias towards class or samples with more cells, we scale the parameter *C* by multiplying with a cell factor *p_i_* in SVM loss function. So, a high value of *p_i_* means less regularization for the cell and a higher incentive for the SVM to classify it properly.

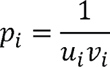

Where *u_i_* is the class frequency of cell *k*, *v_i_* is the sample frequency of cell *i*

#### Stable Gene Selection

To enhance the robustness of gene selection, we incorporate a stable gene selection mode into scPanel. This approach involves the iterative downsampling of the training dataset without replacement, creating multiple subsets. Each subset is then subjected to the scPanel’s minimal gene panel selection algorithm. Genes frequently selected across these iterations are identified as the stable gene panel. We show this procedure in analyzing the severe COVID-19 dataset (wilk2020covid)^11^, where we apply downsampling rates of 70%, 80%, and 90% to the training data, each iterating 20 times. Genes exhibiting a selection frequency exceeding 50% are designated as stable genes. We allow users to specify the downsampling rates, number of iterations, and selection frequency threshold in scPanel.

#### Model training and Patient-level classification

Given the selected most responsive cell type c * and minimal gene panel *R*. We subset the gene expression matrix *D* and class label *Y* into:

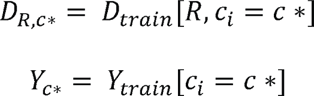

We use *D_R,c*_* and *Y_C*_* as inputs to train 5 different classifiers including:

Logistic Regression (LR): *f_LR_*

Support Vector Machine (SVM): *f_SVM_*

Random Forest (RF): *f_RF_*

k-Nearest Neighbors (kNN): *f_kNN_*

Graph Attention Network (GAT): *f_GAT_*

During the testing phase, for each cell, we computed the final prediction outcome by taking the median of the prediction probabilities generated by the ensemble of the 5 classifiers.

- For LR it naturally outputs probabilities:

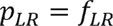
- For SVM, the prediction probability of each cell is estimated using Platt’s method [62] as follows:

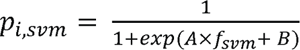 where A and B are parameters (estimated by maximum likelihood) of sigmoid link function that converts the output *f_SVM_* from the SVM into a probability.
- For RF, the prediction probability of each cell is estimated from a collection of decision trees as follows:

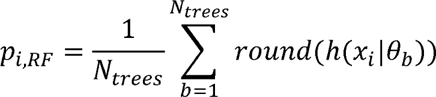 Where *h*(*x*|θ*_b_*) denote decision tree, each parameterised by *θ_b_* by training on the *b*th bootstrap data sample and a subset of features randomly sampled. The *round()* function converts the probability value to the nearest integer to either 0 or 1. We estimate the probabilities by making a class prediction for each tree, and counting the fraction of trees that vote for a certain class.
- For kNN, it use the proportion of the k-nearest neighbors that belong to a certain class as the probability:

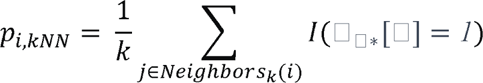 where *Neighbors_k_*(i) denotes the set of indices of the k-nearest neighbors of the cell. The *I*(.) function converts the true or false to either 1 or 0.
- For GAT, it outputs probabilities by considering the features of nodes (cells) and the graph structure. For a cell i, the probability of belonging to class 1 can be represented as:

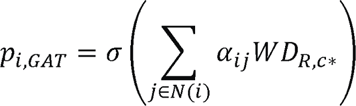 where *N*(*i*) is the set of neighbors of cell i, *α_ij_* are the attention coefficients, *w* is a weight matrix, and < is the sigmoid activation function.

After obtaining the prediction probabilities from each of the classifiers for a given cell, median of these probabilities is calculated as final prediction outcome for the given cell:

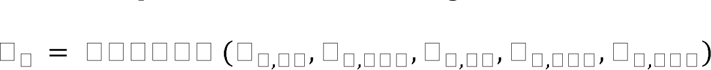

Subsequently, for each sample, we calculated the Area Under the Receiver Operating Characteristic (AUROC) curve. This was accomplished by first ranking the cells within the sample based on their median prediction probabilities. These ranks served as the x-axis, whereas the corresponding median probabilities were plotted on the y-axis. The area under this curve is computed as AUROC for each sample, serving as the measure of the patient-level classification performance. Sample with *AUROC*≥ 0.5 is predicted as 1, otherwise the sample is predicted as 0.

In order to evaluate the statistical significance of the sample-level predictions, we assessed whether the AUROC was significantly different from that of a random classifier, having an expected AUROC of 0.5. To achieve this, we employed a non-parametric bootstrap approach. As described in the last paragraph, we denoted the sample-level AUROC of the predictive model computed on the testing data as *A_sample_*. Under the null hypothesis that the model is no better than random guessing, we set *H*_0_:*A*=0.5. Subsequently, we generated *N* bootstrap samples {*D_1_*,*D*_2_,…,*D_N_*} from the original testing data. For each bootstrap sample *D_i_*, the AUROC *α_i_* was computed, resulting in a distribution of AUROC values under the null hypothesis. The empirical p-value was calculated to quantify the significance of *A_sample_*. It was defined as the proportion of bootstrap AUROC values that were as extreme as or more extreme than *A_sample_* represented as:

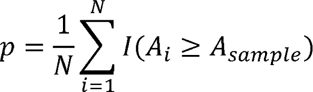

where *l* is an indicator function, yielding 1 if *A_i_* ≥*A_sample_* and 0 otherwise. A small empirical p-value indicates that the observed AUROC is significantly different from what would be expected under the null hypothesis, hence suggesting that the predictive model performs better than random guessing.

#### Data preprocessing

Gene expression matrices were first columns normalized and log transformed with *NormalizeData()* in the Seurat R package. Genes that are expressed in at least one cell are retained in the data.

#### Cross-batch prediction

Testing data was integrated with training data using *Seurat V3 CCA* method^43^. *SelectIntegrationFeatures()* was used to find the top 2000 highly variable genes shared by both data as integration features. *FindIntegrationAnchors()* was used to find anchor features with parameter *n_PC = 20*. *IntegrateData()* was used for integration with parameters *n_PC = 20*, *k_weight = 100*.

#### Benchmarking with differentially expressed genes, ActiveSVM and COMET

We compared the scPanel gene selection procedures: (*i*) differentially expressed genes (DE genes)^43^, (*ii*) ActiveSVM^8^ and (*iii*) COMET^9^. For (*i*), *FindMarkers()* from the Seurat R package was used to identify differentially expressed genes using a Wilcoxon Rank Sum test. Genes were ranked by absolute values of average log fold changes. For (*ii*), Min-complexity mode was run with parameters *num_samples = 100, init_samples= 200, balance=True* for COVID-19 dataset (wilk2020covid) and *num_samples = 200, init_samples = 400* for scleroderma dataset (gur2022ssc); Min-cell mode was run with parameters *num_samples = 100, init_samples = 200, balance=True* for COVID-19 dataset (wilk2020covid) and *num_samples = 200, init_samples = 400* for scleroderma dataset (gur2022ssc). (*iii*) COMET was run with default parameters.

The comparisons were done using 1) top 4 genes identified by each method, a constraint imposed due to COMET’s limitation of exhaustively searching a maximum of four gene combinations; 2) the top number of genes automatically decided by scPanel, as other methods rely on users to arbitrarily specify the number of genes.

Patient-level AUROC is used as the benchmarking metric. Specifically, the same training data was input to these methods to select genes. The patient-level classification model training and testing were uniformly conducted using the same parameters in scPanel. Patient-level AUROC is computed for each sample relative to its specific class in the testing data.

### Empirical Power Analysis

The empirical power of scPanel was estimated with respect to the number of cells and the number of samples (scRNA-seq datasets), as follows. For cell-level power analysis, we downsampled the training set by cell to different proportions, each repeated for 20 times. The downsampling was done without replacement and stratified by sample and cell type. Sample-level power analysis was done similarly by downsampling the training set by sample to different proportions, each repeated for 20 times. The testing set remains the same. The downsampled training set is used to run scPanel for cell type selection, gene selection and patient-level classification.

#### Assessment of the elapsed computational time and peak memory usage

All tasks were performed on a research High Performance Computer (HPC) system with AMD EPYC 7763 64-Core Processors. 50 CPUs and 256 Gb total memory are allocated. The elapsed computational time was evaluated by the function ‘time()’ from the *time* python module; timings for each method include all pre-processing steps. The usage of peak memory was monitored by the function ‘memory_usage()’ from *memory_profiler* python module. Random seeds were fixed for all steps in the scPanel to ensure reproducibility.

## Supplementary materials

Supplementary Figure 1 - 7

Supplementary Table 1 - 2

## Data availability

All of the data used in the paper have been previously published. COVID-19 single cell RNA-seq data (wilk2020covid, su2020covid, lee2020cov_flu) are downloaded from COVID-19 Atlas (https://atlas.fredhutch.org/fredhutch/covid). Scleroderma single cell RNA-seq data (gur2022ssc) are downloaded from NCBI Gene Expression Omnibus (GEO) with accession number GEO: GSE195452.

## Code availability

Our method is integrated as an installable Python package called scPanel. The installation instructions and user guidance are shown at https://test.pypi.org/project/scPanel/. The source code of scPanel and demo examples will be publicly available on GitHub upon publication.

## Ethics approval and consent to participate

Not applicable.

## Authors’ contributions

Conceptualization, Yi Xie with input from John Ouyang and Enrico Petretto; Data curation and analysis, Yi Xie; Formal analysis, Yi Xie; Visualization, Yi Xie; Writing – original draft, Yi Xie, John Ouyang; Writing – review & editing, Yi Xie, John Ouyang, Enrico Petretto, Jianfei Yang; Project supervision, Enrico Petretto, John Ouyang. All authors have read and agreed to the published version of the manuscript.

