## Supplementary Figures for "scPanel: A tool for automatic identification of sparse gene panels for generalizable patient classification using scRNA-seq datasets"

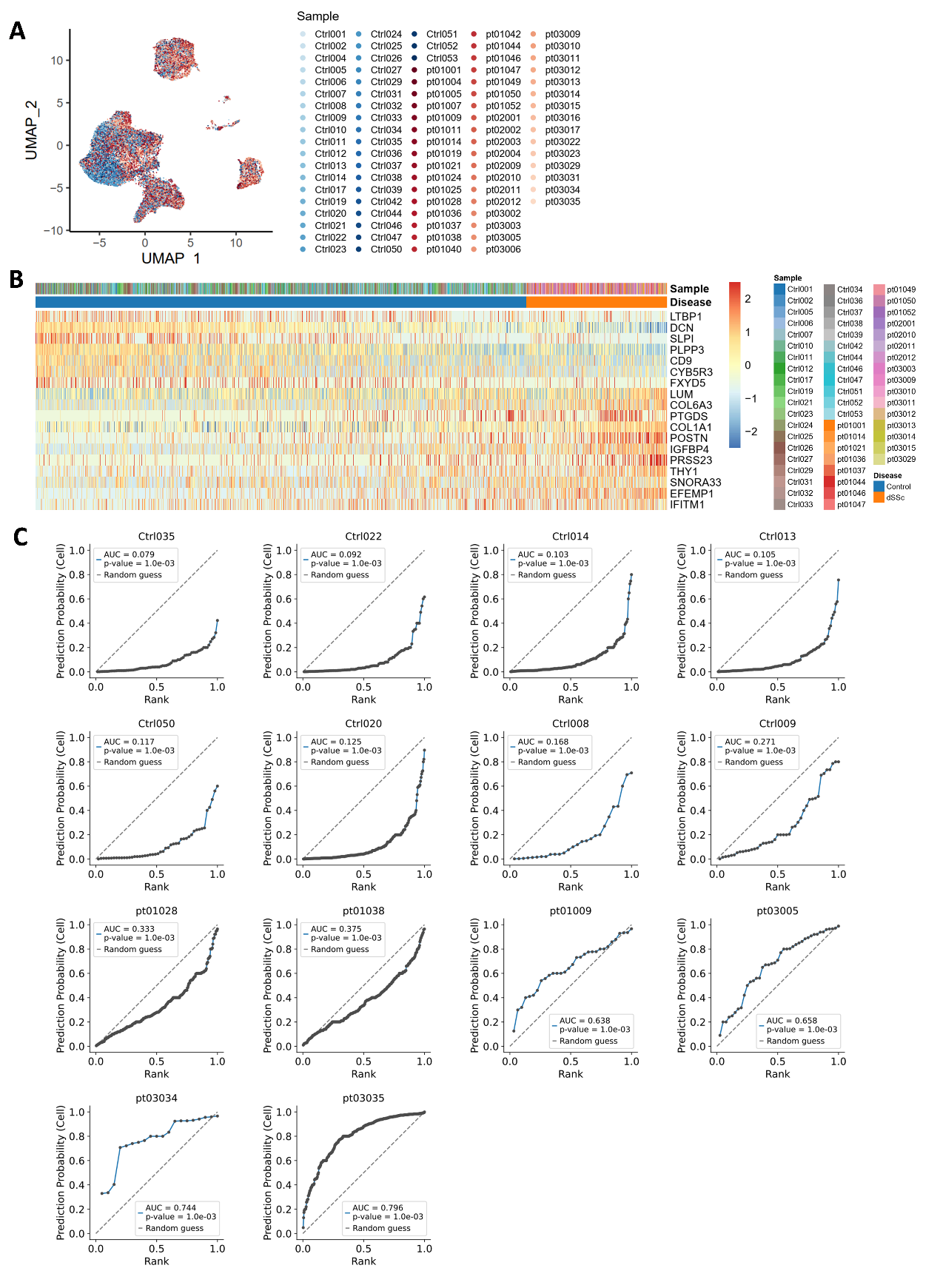

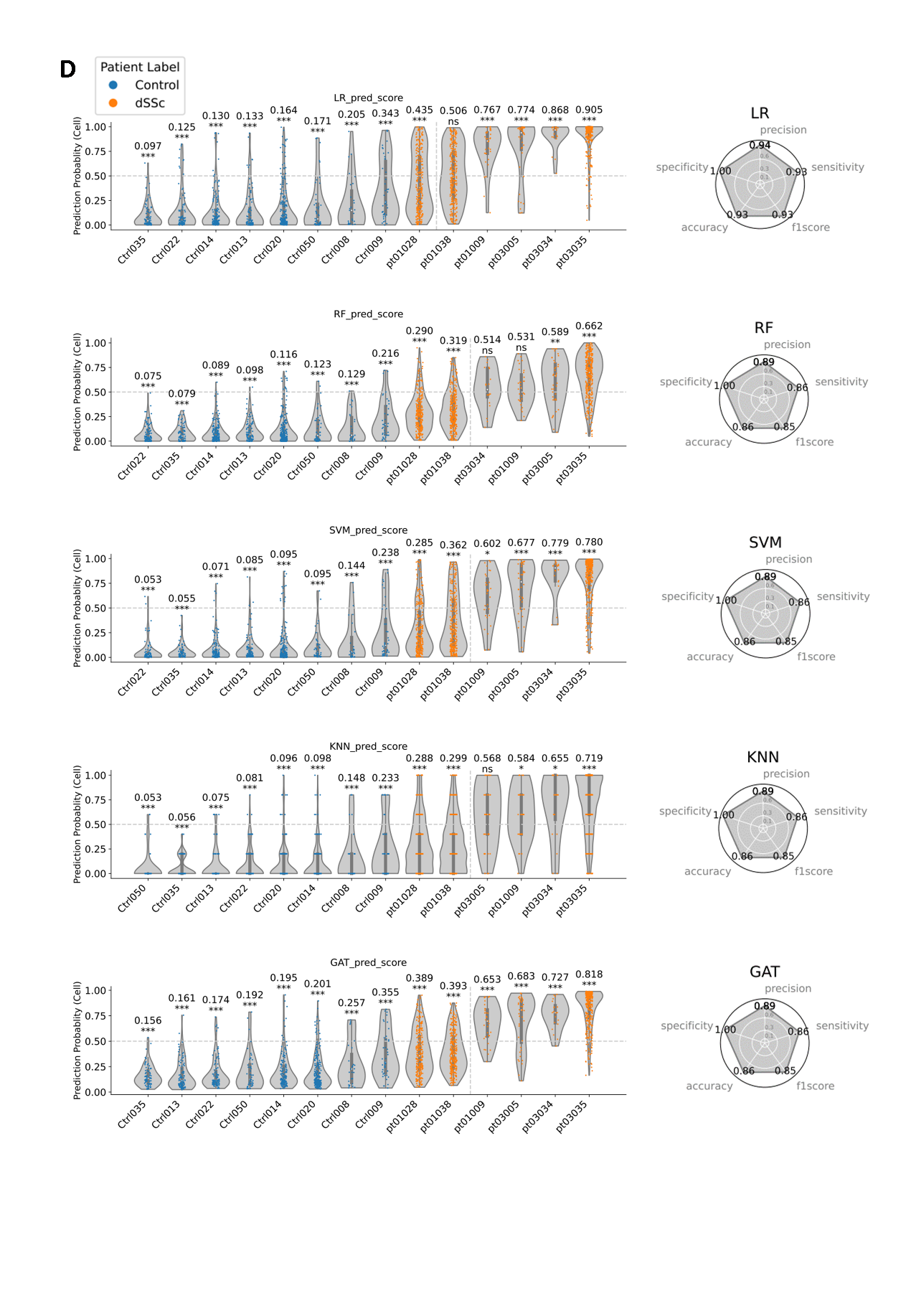

**Supplementary Figure 1. Using scPanel to classify diffuse cutaneous systemic sclerosis (dSSc) patients**

A) UMAP of input dataset colored by samples. B) Heatmap of gene expression of selected genes in the training set. Values are z-score of normalized gene expression truncated at ±2.5. C) Receiver operating characteristic curve (ROC) for prediction of each sample. D) Violin plot of patient-level prediction results in each of the 5 classifiers. Violins are colored by the label of patients. Y-axis represents cell-level prediction probability. Value on top of each violin represents sample-level AUC score with p-value (* p<.05; ** p<.01; *** p<.001). The vertical line indicates the patient-level prediction threshold (AUC = 0.5). Patients at the right side of this line are predicted as dSSc patients. Accuracy, precision, sensitivity and recall are computed to evaluate the overall patient classification performance.

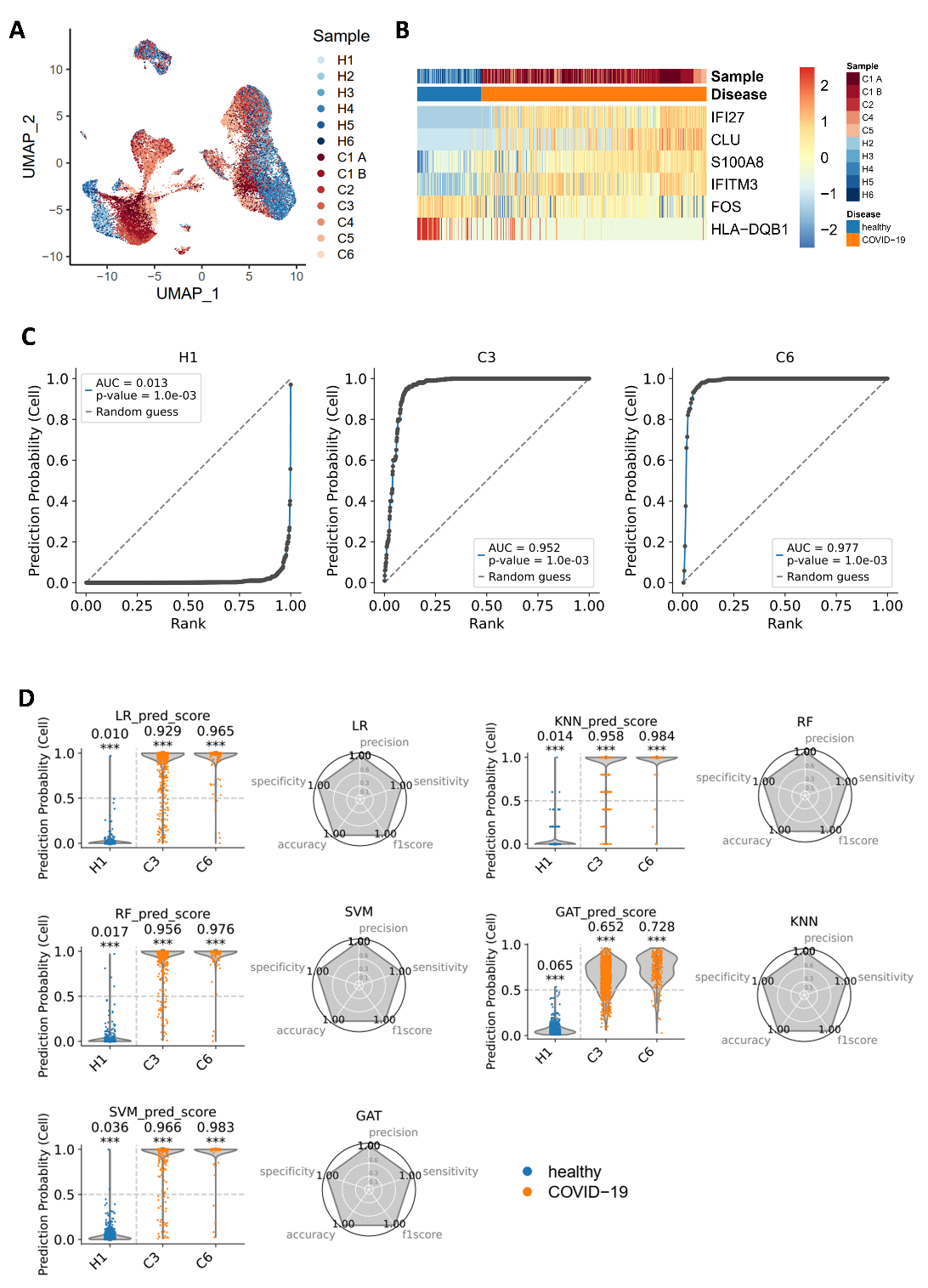

**Supplementary Figure 2. Using scPanel to classify severe SARS-CoV-2 (COVID-19) patients**

A) UMAP of input dataset colored by samples. B) Heatmap of gene expression of selected genes in the training set. Values are z-score of normalized gene expression truncated at ±2.5. C) Receiver operating characteristic curve (ROC) for prediction of each sample. D) Violin plot of patient-level prediction results in each of the 5 classifiers. Violins are colored by the label of patients. Y-axis represents cell-level prediction probability. Value on top of each violin represents sample-level AUC score with p-value (* p<.05; ** p<.01; *** p<.001). The vertical line indicates the patient-level prediction threshold (AUC = 0.5). Patients at the right side of this line are predicted to be severe COVID-19 patients. Accuracy, precision, sensitivity and recall are computed to evaluate the overall patient classification performance.

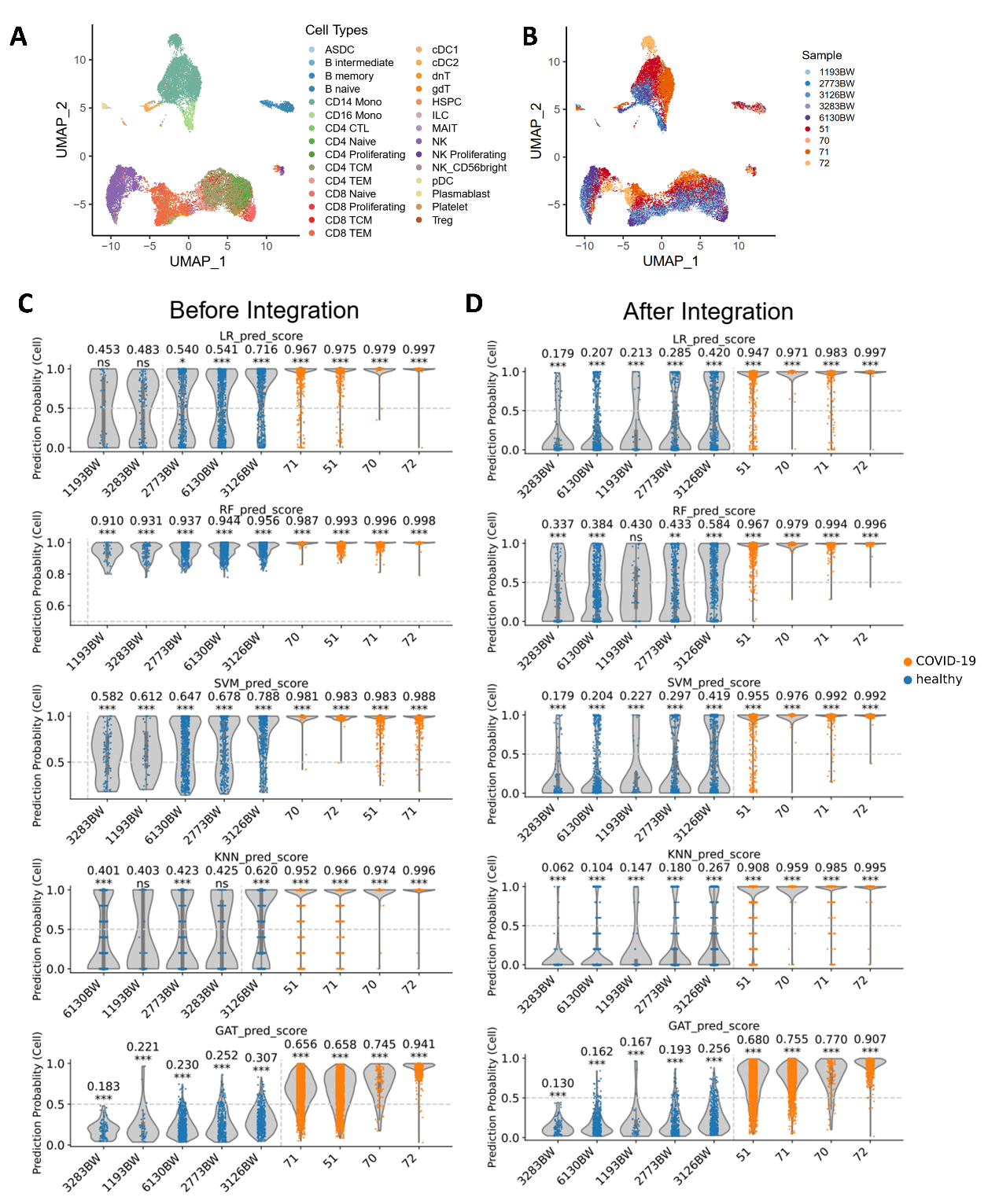

**Supplementary Figure 3. Using scPanel to predict cross-batch severe SARS-CoV-2 (COVID-19) patients**

A, B) UMAP of cross-batch testing data (su2020covid) colored by A) cell types and B) samples. C, D) Violin plot of patient-level prediction results in each of the 5 classifiers in testing data (su2020covid) C) without data integration and D) after data integration. Violins are colored by the label of patients. Y-axis represents cell-level prediction probability. Value on top of each violin represents sample-level AUC score with p-value (* p<.05; ** p<.01; *** p<.001). The vertical line indicates the patient-level prediction threshold (AUC = 0.5). Patients at the right side of this line are predicted to be severe COVID-19 patients.

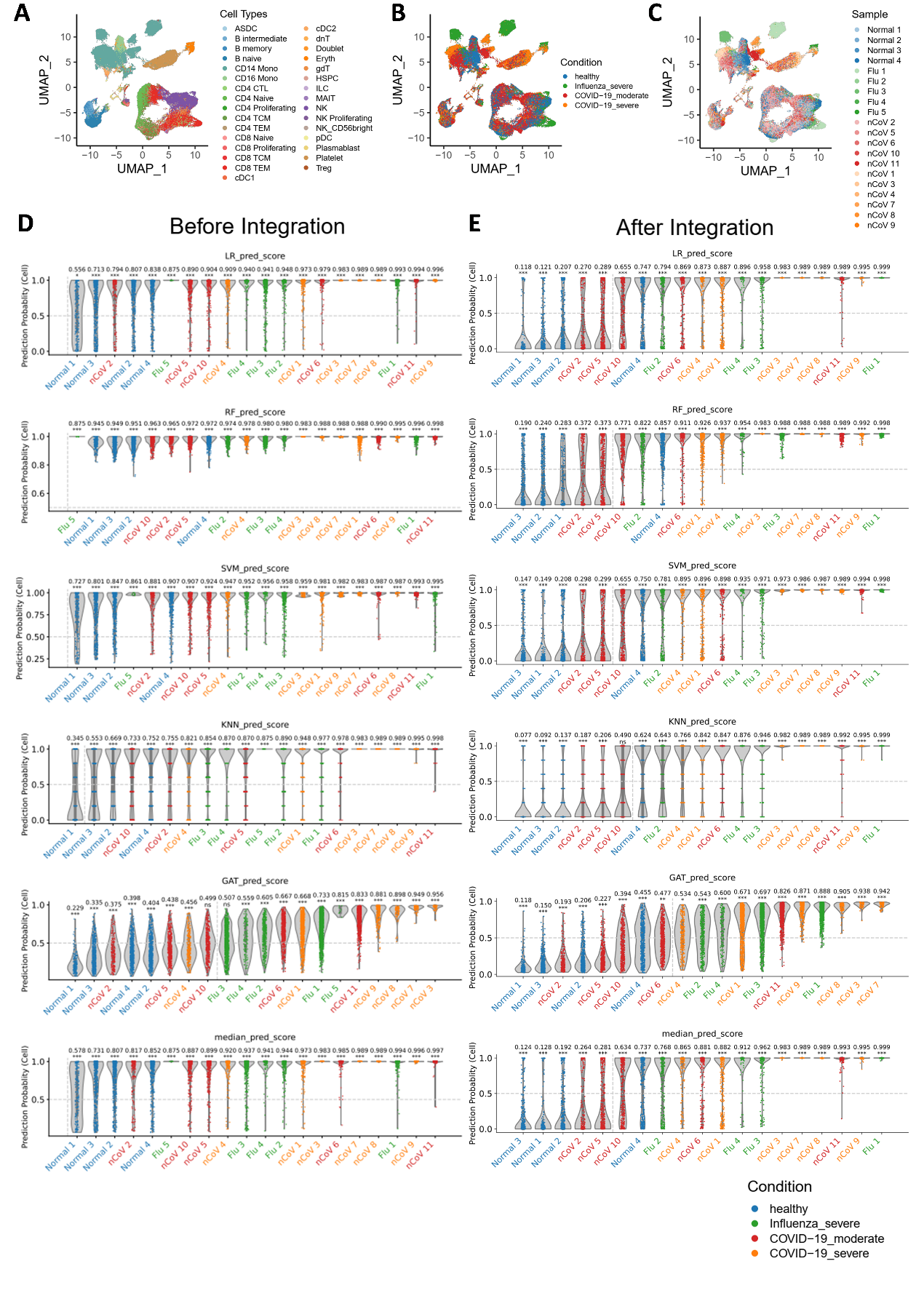

**Supplementary Figure 4. Specificity of severe SARS-CoV-2 (COVID-19) biomarkers derived from wilk2020covid dataset by scPanel**

A-C) UMAP of cross-batch testing data (lee2020cov_flu) colored by A) cell types, B) conditions and C) samples. D, E) Violin plot of patient-level prediction results in each of the 5 classifiers and median score in testing data (su2020covid) D) without data integration and E) after data integration. Violins are colored by the label of patients. Y-axis represents cell-level prediction probability. Value on top of each violin represents sample-level AUC score with p-value (* p<.05; ** p<.01; *** p<.001). The vertical line indicates patient-level prediction threshold (AUC = 0.5). Patients at the right side of this line are predicted to be severe COVID-19 patients.

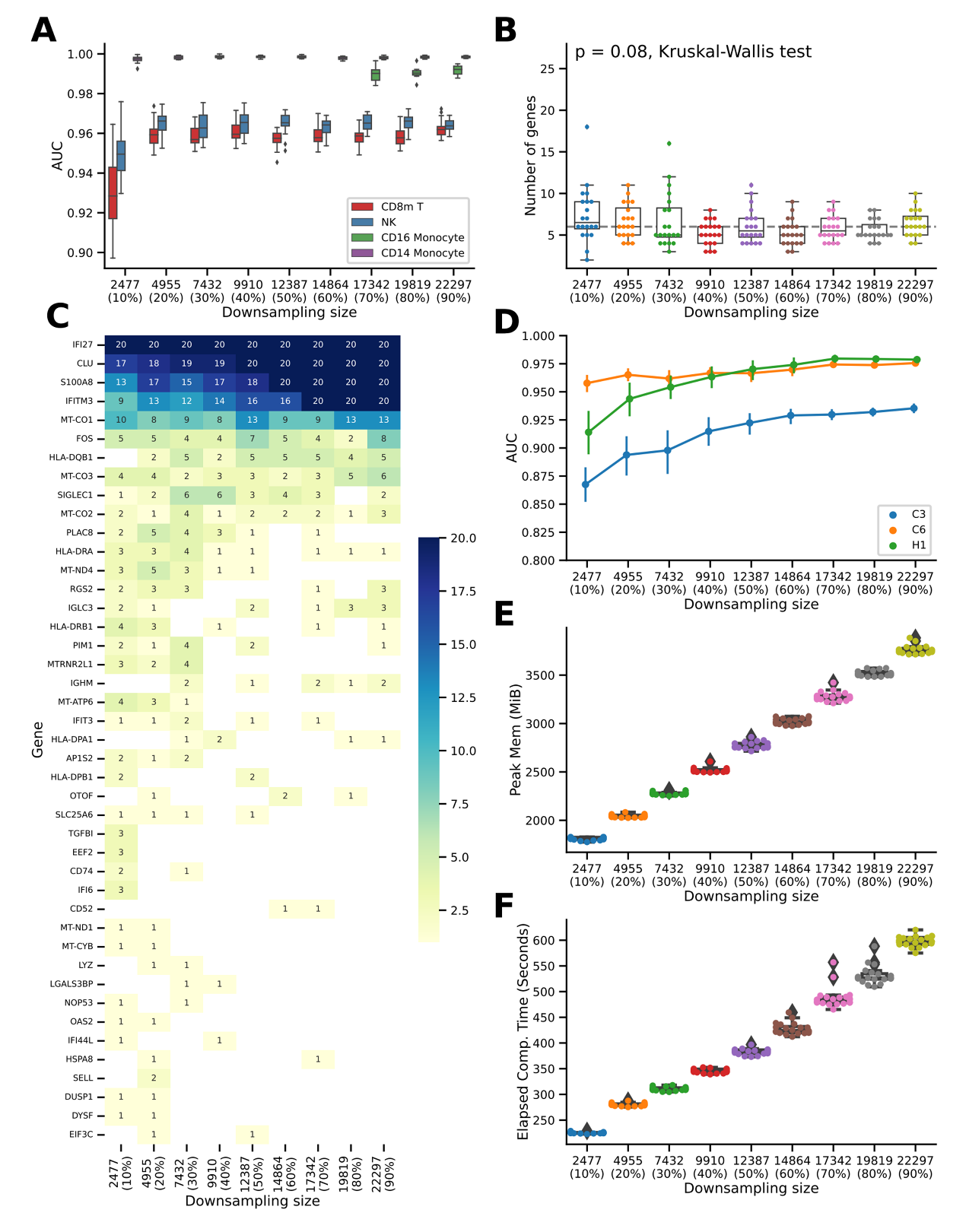

**Supplementary Figure 5. Cell-level power analysis of scPanel in severe COVID-19**

A) AUC scores for the top four most responsive cell types, quantifying their response to severe COVID-19 perturbation across various downsampling sizes (number of cells). B) Number of genes selected across various downsampling sizes for CD14+ monocytes. C) Gene selection frequency across various downsampling sizes for CD14+ monocytes. Each value represents the number of times that a specific gene is selected out of 20 iterations. D) AUC scores of patient-level predictions across various downsampling sizes. E) Peak memory usage (mebibyte, MiB) for executing across various downsampling sizes. F) Elapsed computational time (seconds) for executing scPanel across various downsampling sizes.

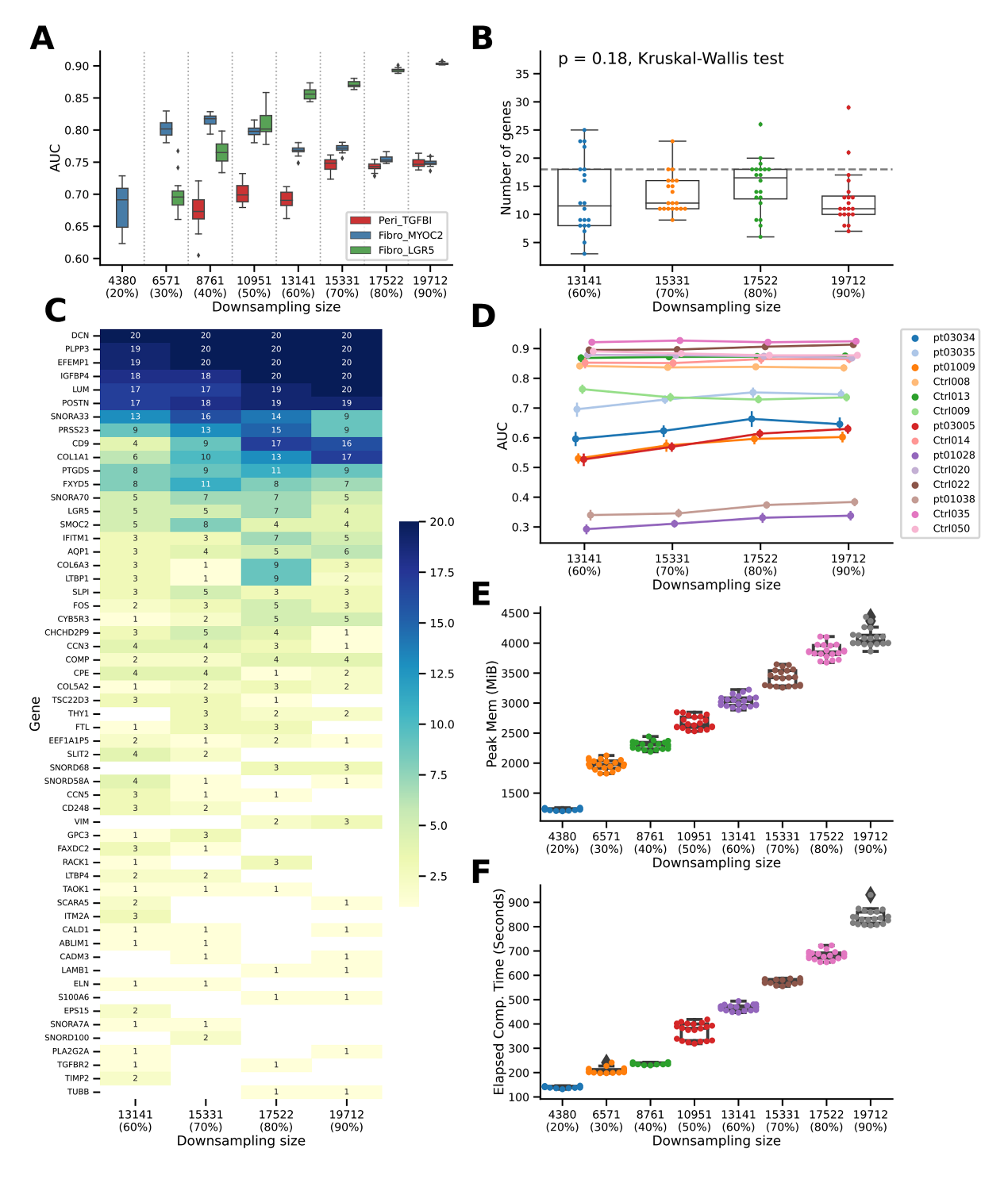

**Supplementary Figure 6. Cell-level power analysis of scPanel in dSSc**

A) AUC scores for the top four most responsive cell types, quantifying their response to dSSc perturbation across various downsampling sizes (number of cells). B) Number of genes selected across various downsampling sizes for LGR5+ fibroblasts. C) Gene selection frequency across various downsampling sizes for LGR5+ fibroblasts. Each value represents the number of times that a specific gene is selected out of 20 iterations. D) AUC scores of patient-level predictions across various downsampling sizes. E) Peak memory usage (mebibyte, MiB) for executing across various downsampling sizes. F) Elapsed computational time (seconds) for executing scPanel across various downsampling sizes.

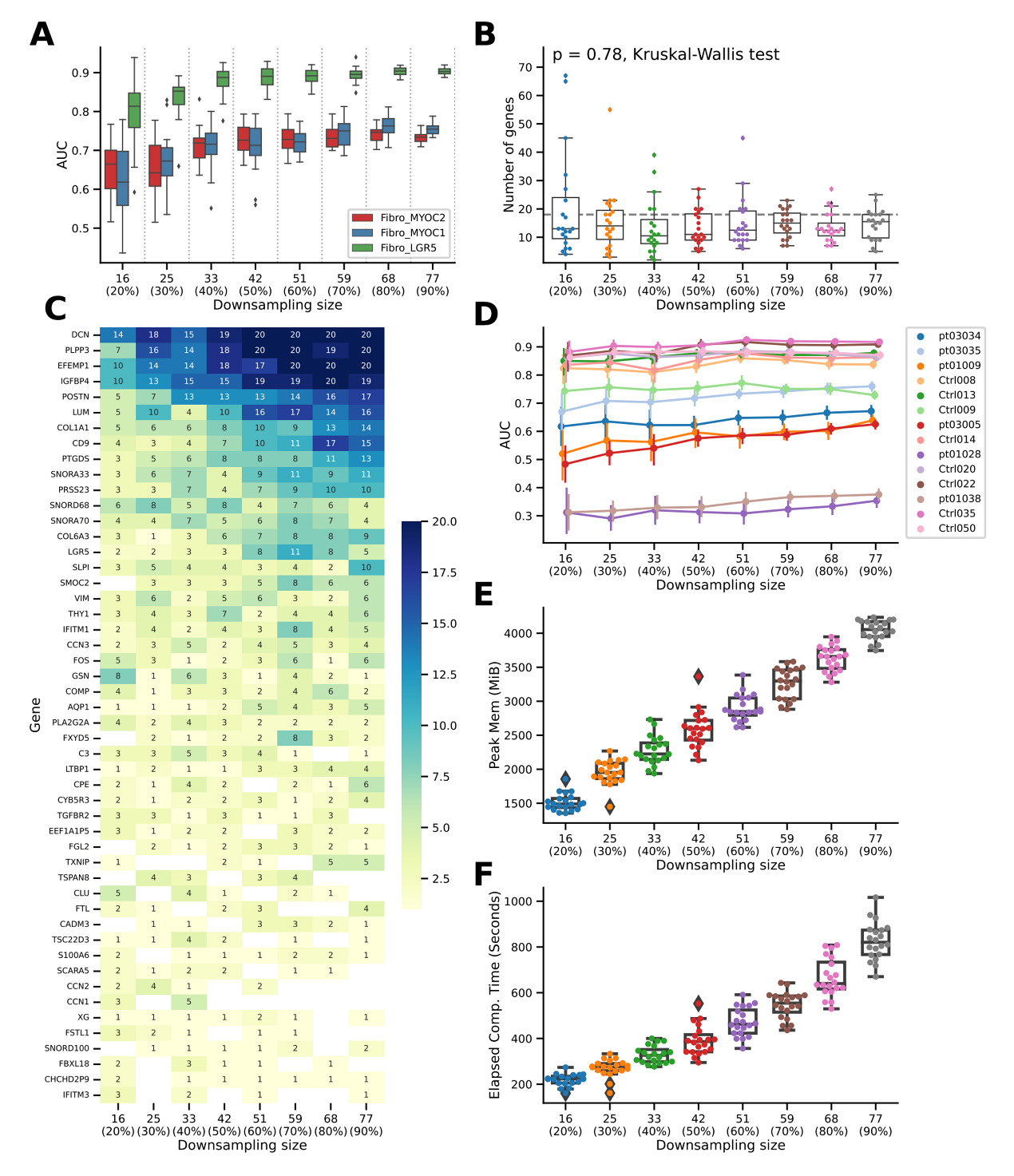

**Supplementary Figure 7. Sample-level power analysis of scPanel in dSSc**

A) AUC scores for the top four most responsive cell types, quantifying their response to severe COVID-19 perturbation across various downsampling sizes (number of samples). B) Number of genes selected across various downsampling sizes for LGR5+ fibroblasts. C) Gene selection frequency across various downsampling sizes for LGR5+ fibroblasts. Each value represents the number of times that a specific gene is selected out of 20 iterations. D) AUC scores of patient-level predictions across various downsampling sizes. E) Peak memory usage (mebibyte, MiB) for executing across various downsampling sizes. F) Elapsed computational time (seconds) for executing scPanel across various downsampling sizes.
